# Genetic admixture history and forensic characteristics of Tibeto-Burman-speaking Qiang people explored via the newly developed Y-STR panel and genome-wide SNP data

**DOI:** 10.1101/2022.04.13.488250

**Authors:** Guanglin He, Atif Adnan, Mengge Wang, Wedad Saeed Al-Qahtani, Fatmah Ahmed Safhi, Hui-Yuan Yeh, Sibte Hadi, Chuan-Chao Wanag, Chao Liu, Jun Yao

## Abstract

Fine-scale patterns of population genetic structure and diversity of ethnolinguistically diverse populations are important for biogeographical ancestry inference, kinship testing and also for the development and validation of new kits focused on forensic personal identification. Analyses focused on forensic markers and genome-wide SNP data can provide new insights into the origin, admixture processes and forensic characteristics of targeted populations. Qiang people with a large sample size among Tibeto-Burman-speaking populations widely reside in the middle latitude of the Tibetan Plateau. However, their genetic structure and forensic features have remained uncharacterized due to the paucity of comprehensive genetic analyses. Here, we first developed and validated the AGCU-Y30 Y-STR panel, which contains slowly and moderately mutating Y-STRs, and then we conducted comprehensive population genetic analyses based on Y-STRs and genome-wide SNPs to explore the admixture history of Qiang people and their neighbours. The validated results of this panel showed that the new Y-STR kit was sensitive and robust enough for forensic applications. Haplotype diversity (HD) ranging from 0.9932 to 0.9996 and allelic frequencies ranging from 0.001946 to 0.8326 in 514 Qiang people demonstrated that all included markers were highly polymorphic in Tibeto-Burman people. Population genetic analyses based on Y-STRs (R_ST_, F_ST_, MDS, NJ, PCA and MJNs) revealed that the Qiang people harboured a paternally close relationship with lowland Tibetan-Yi corridor populations. Furthermore, we made a comprehensive population admixture analysis among Eurasian modern and ancient populations based on the shared alleles. We determined that the Qiang people were a genetically admixed population and showed the closest relationship with Tibetan and Neolithic Yellow River farmers. Admixture modelling showed that Qiang people shared the primary ancestry with Tibetan and was derived from North China, supporting the hypothesis of common origin between Tibetan and Qiang people.

## 1 INTRODUCTION

East Asia has more than eight language families or groups, including Tungusic, Mongolic, Turkic, Sino-Tibetan, Austronesian, Austroasiatic, Tai-Kadai and Hmong-Mein, and over hundreds of ethnic groups and harboured complex patterns of genetic diversity. Recent comprehensive genetic analyses have shown that population genetic analyses based on geographically and ethnically different populations could provide new insights into the formation of modern populations, especially in some regions with rich ethnolinguistic diversity [1,2]. Recent advances in population genomic studies have provided an updated landscape of the genetic basis of East Asians [3]. ChinaMap project based on the whole-genome sequences of over 10K people showed significant genetic differentiation between Han Chinese and other minorities and found seven substructured groups of Han Chinese [4]. Genome-wide SNP data from northern, central, southern, and western China also illuminated the different ancestral source composition and admixture weight in the formation of geographically diverse modern Hans [5–11]. Furthermore, recent ancient DNA studies provided new insights into the peopling of the Tibetan Plateau and modern Sino-Tibetan people [3,8,11,12]. Ancient genomes from Nepal showed their genetic connection to East Asians rather than South Asians, ancient northern East Asian genomes from Gansu, Henan and Shaanxi provinces also showed the close genetic relationship between millet farmers and modern Tibetans [3]. These genetic findings showed complex patterns of genetic diversity and the population structure of East Asia. Similar patterns of the consistent association between African genetic diversity and four language families (Afro-Asiatic, Nilo-Saharian, Niger-Saharian (Niger-Congo), and Khoisan) were also identified in East Asia [13]. Altaic-speaking populations in North China showed predominant ancestry related to ancient Mongolian ancestry and western Eurasian ancestry [14,15]. Other populations from Sino-Tibetan language families in central and southern East Asia harboured more primary ancestry related to ancient Yellow River Basin and Yangtze River Basin ancestry [9,10,16]. These population stratifications provided a new opportunity for the development and validation of new forensic kits focused on the fine-scale genetic background of East Asians with different genetic markers and higher resolution. This type of the kit could provide for a more comprehensive genetic landscape reconstruction of East Asians.

Genetic markers localized in the non-recombining regions of Y-chromosome have played an important role in forensic identification, kinship testing and molecular anthropology. The initial peopling history of East Asians has been explored via Y-chromosome markers, such as the demic diffusion of Han Chinese rather than the cultural diffusion [17]. Besides the function of population history reconstructions, Y-STRs are widely used for characterizing male contributions to male-female mixtures in forensic genetics, particularly in sexual assault cases or familial searches such as kinship casework involving male offspring, especially in deficient paternity cases where the putative father was unavailable [18–21]. Currently, many Y-STR commercial kits are available, such as Yfiler (ThermoFisher Scientific) [22], Yfiler Plus (ThermoFisher Scientific) [23], GoldenEye 20Y (PEOPLESPOT R&D, China), GoldenEye 26Y (PEOPLESPOT R&D, China), DNATyper Y26 (Second Institute of the Ministry of Public Security) [24], Powerplex Y23 (Promega) [25], and AGCU Y24 (China-Germany United) [26]. These kits usually contain various numbers of Y-STRs ranging from 17 to 27 and among these 3 to 7 rapidly mutating Y-STRs have been included [27]. Several studies have demonstrated that Y-STR haplotype diversity and male lineage differentiation can be improved by adding additional carefully chosen Y-STRs[28]. Rapidly mutating Y-STRs with mutation rates > 1 × 10^-2^ have been used for the differentiation of closely related male individuals and can help individualize them due to their high discrimination power (DP) [29–31]. However, when it comes to familial searching, these Y-STRs are not conclusive. Keep this in mind, the AGCU has introduced a new 6-dye system that can co-amplify 30 Y-STRs: DYS392, DYS389I, DYS447, DYS389II, DYS438, DYS527a/b, DYS522 labeled with FAM, DYS391, DYS456, DYS19, DYS388, DYS448, DYS385a/b labeled with HEX, DYS481, DYS437, DYS390, DYS460, DYS533, DYS458 labeled with TAMRA, DYS393 Y_GATA_H4, DYS439, DYS635, DYS444, DYS643 labeled with ROX while DYS549, DYS557, DYS520 labeled with VIG dye (**Supplementary Figure 1**). These 30 Y-STRs have low mutation rates (1 × 10^-3^ to 1 × 10^-5^) which make them good candidates for family lineage reconstruction and are helpful for the construction of Y-STR DNA database, forensic casework and kinship cases because of having reduced risk of homogeneity and heterogeneity. The AGCU Database Y30 kit’s accuracy, stability, sensitivity, specificity, resistance to inhibition, and mixture sample analysis were tested in the Qiang population from Beichuan, Sichuan, China.

Sino-Tibetan-speaking populations were the largest population group in Asia, which were separated into Tibeto-Burman speakers residing in the Tibetan Plateau and Sinitic-speaking people in the low altitude regions of East Asia. The Qiang people are an ancient group in mainland China, which is associated with the spread of modern Sino-Tibetan language and cultures. Their presence has been noticed on the Oracle bone which is known as a “living fossil” in the history of the Chinese nation and is almost 3000 years old. Many of these people in past were designated as “Qiang” and later on these were also influenced by Chinese culture and amalgamated with the Han population during Ming and Qing dynasties, these people were again reclassified and non-Han people who were living in the upper valley of Min river and Beichuan area are called Qiang [32]. Qiang people speak their own language of Qiangic, a subfamily of the Tibeto-Burman languages. They are one of 55 minority ethnic groups in China. All minorities are 8.49% of the total Chinese populations and Qiang are 0.23% of minority populations with a population size of 310000 and one-third of their population is located in the Beichuan Qiang Autonomous County [33]. Beichuan Qiang Autonomous County is associated with Mianyang City, Sichuan Province, China and is the only Qiang autonomous county in China.

Previous genetic analysis has mainly focused on the fine-scale population genetic structure of Han Chinese and Tibetan people using genome-wide SNP data, however, this important Qiang population in this language family keeps uncharacterized until now [11,34]. There is limited knowledge of the genetic diversity and fine-scale admixture structure of Qiang people and their genetic origin and detailed evolutionary history reconstructed based on the genome-wide SNP data also remains uncharacterized. Thus here, we first validated a Y-STR genotyping panel to obtain specific Y-STR data and also conducted a genome-wide SNP study. This was aimed to determine the forensic efficiency and application of the new-developed AGCU Database Y30 kit, study the genetic relationships between Qiang people and their neighbours based on the low-moderate mutated Y-STRs and to determine the fine-scale genetic structure, admixture history and development of modern Qiang people based on the genome-wide SNP data.

## 2 MATERIALS AND METHODS

### 2.1 Samples preparation

Control DNA 9948, 007 (male DNA) and 9947A (female DNA) were used for sensitivity, precision evaluation, inhibitor study, and DNA mixture studies. Blood samples were collected from a total of 534 unrelated healthy individuals from Qiang ethnic group (514 male and 20 female) from Beichuan Qiang Autonomous County, Mianyang City, Sichuan Province, China (**Figure 1**). All participants gave their informed consent either orally and with thumbprints (in case they could not write) or in writing after the study aims and procedures were carefully explained to them in their language. The study was approved by the ethical review board (Dated 20^th^ March 2019 with approval reference no. 2019-86-P) of the China Medical University, and Xiamen University (XDYX2019009), People’s Republic of China. Whole blood samples of pigs, cattle, sheep, chickens, ducks, fish, rats and rabbits were purchased from Wuxi Zoological Garden (Wuxi, Jiangsu, China). All blood samples were stored at −20 °C before DNA extraction. DNA was isolated from blood using ReliaPrep^™^ Blood gDNA Miniprep System (Promega, Madison, USA) according to the manufacturer’s instructions. DNA quantitation for all samples was determined using a NanoDrop spectrophotometer (Thermo Scientific, Wilmington DE, USA) and the final concentration of DNA was diluted to 1 to 2ng/μl. Moreover, 9948 was also used as positive/ control sample in all PCR batches.

**Figure 1.**
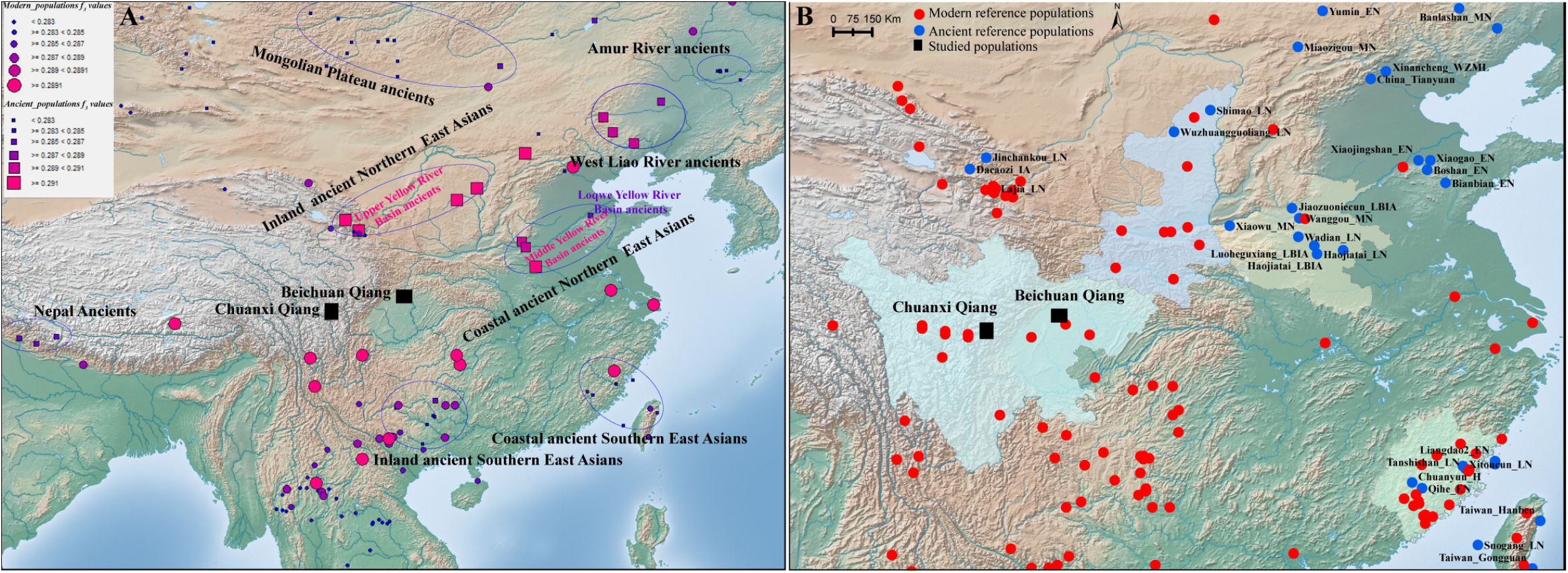
Geographical positions of our studied populations and reference populations. (**A**) The genetic affinity between Chuanxi Qiang and reference populations from HO dataset. The color and size of the circle showed the genetic affinity between Qiang people and modern reference populations and the color and size of the square denoted the genetic affinity with ancient reference populations. (B) The geographical position of studied populations and reference populations from the Illumina dataset.

### 2.2 Reference dataset

Y-STR data from different populations were collected from the YHRD database. Genome-wide SNP data of Qiang people developed by us was used for this work and then merged with other modern and ancient genomes from China, Nepal, Southeast Asia, Japan, Mongolia and Siberia collected from the Allen Ancient DNA Resource (AADR) (https://reich.hms.harvard.edu/allen-ancient-dna-resource-aadr-downloadable-genotypes-present-day-and-ancient-dna-data).

### 2.3 Validation framework

#### 2.3.1 PCR amplification and electrophoresis

PCR amplification was performed in a GeneAmp^®^PCR System 9700 thermal cycler (Thermofisher Scientific, Foster City, CA, USA) on a gold-plated silver block. After optimization, the final reaction system was performed in a 25 μL volume containing 10 μL of master mix, 5 μL of primer set, 1 μL of 5U/μL Taq DNA polymerase (AGCU ScienTech Incorporation, Wuxi, Jiangsu, China), and 0.5–2 ng of gDNA under the following PCR conditions: initial denaturation of 95 °C for 2 min, 30 cycles of denaturation, annealing and extension step at 94 °C for 30 s, 59 °C for 60 s, 72 °C for 60 s, followed by a final extension step at 60 °C for 30 min and a final soak at 4 °C. Amplification products were analyzed regarding AGCU Marker SIZ-500 internal size standard (AGCU Incorporation, Wuxi, China) and AGCU Database Y30 kit Allelic Ladder using an ABI 3500 genetic analyzer (Thermofisher Scientific, Foster City, CA, USA) with the POP-4^™^ polymer (Thermofisher Scientific, Foster City, CA, USA) according to AGCU Database Y30 kit standard protocol. Injection time of 10 s, injection and run voltages 3 KV and run time 2,500 s were used for all electrophoresis runs on 3500 genetic analyzer. Gene-Mapper Software version 4.0 (Thermofisher Scientific, Foster City, CA, USA) was used for genotype assignment.

#### 2.3.2 Sensitivity test

Serial dilutions of control DNA 9948 were prepared using dH_2_O (1.0 ng/μL, 0.5 ng/μL, 0.25 ng/μL, 0.125 ng/μL, 0.1 ng/μL, 0.05 ng/μL). These were amplified in triplicate using 25 to 10 μL reaction volume. Amplification processes were performed according to the manufacturer’s instructions.

#### 2.3.3 Species specificity test

1-2.5 ng/μL gDNA of cat, sheep, chicken, duck, rabbit, mice, pig, E. coli was used in a 10 μL reaction volume for each animal and amplified according to the above-mentioned PCR conditions.

#### 2.3.4 Consistency test

We have selected 20 unrelated individuals from the Qiang population and genotyped those with the AGCU Database Y30 kit and the same 20 unrelated individuals from the Qiang population were also genotyped with Yfiler plus kit.

#### 2.3.5 Stability test

To test the stability of the AGCU Database Y30 kit, the components of the kit were repeatedly frozen and thawed 5 to 10 times before amplification while the control group was kept at −20°C. Furthermore, common inhibitors which may be part of biological samples were mixed with control DNA 9948 for the subsequent PCR experiments. The concentrations of used inhibitors were diluted as follows: heme (25, 50, 75, 100 and 125 μM), indigo (10, 12, 14, 16 and 18 μM), humic acid (40, 50, 60, 70 and 80 μg/μL), Ca2+ (0.5, 0.75, 1.0, 1.5 and 2.0 μM), hemoglobin (20, 30, 40, 50 and 60 μM) and EDTA (500, 750, 1000, 1500 and 2000 μM) by Sigma-Aldrich, Saint Louis, MO, USA.

#### 2.3.6 Gender-specific test

1-2 ng/μL of gDNA from 20 unrelated female individuals was amplified in a 10 μL reaction volume according to the PCR conditions of the AGCU Database Y30 kit.

#### 2.3.7 Mixture sample analysis

Manually male(9948)/male(007) mixture samples were prepared with the concentration ratio of 1:1, 1:4, 1:9, 1:19, and 1:29, while male (9948)/female (9947A) were also prepared with the same concentration ratio and then amplified according to the PCR conditions of AGCU Database Y30 kit.

#### 2.3.8 PCR variation of primer concentration, annealing temperature and cycle number testing

We have used 10 μL and 25 μL reaction volume with 1X and 2X primers to detect 1 ng of control DNA (9948) and repeat it three times with annealing temperatures 56°C, 57°C, 58°C, 59°C, 60°C, 61°C, 62°C, 63°C while cycle numbers 29 and 30 were used.

### 2.4 Population and concordance studies based on the non-high mutated STRs

#### 2.4.1 Genotyping of Y-STR data and forensic statistically parameter estimation

514 unrelated individuals from the Qiang population were genotyped with the AGCU Database Y30 kit. Allelic and haplotype frequencies were computed by the direct counting method, and haplotype diversity (HD) was calculated according to:

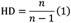

where *n* is the male population size and *p_i_* is the frequency of *i*th haplotype. Discrimination capacity (DC) was calculated as the ratio of unique haplotypes in the samples. Match probabilities (MP) were calculated as Σ P*i*^2^, where P*i* is the frequency of the *i*-th haplotype. We calculated both Rst and Fst values because, in the generalized stepwise mutation model, Rst offers relatively unbiased evaluations of migration rates and times of population divergence, while on other hand F_ST_ tends to show too much population similarity, predominantly when migration rates are low or divergence times are long. Genetic distances were evaluated between reference populations and the Qiang population on overlapping STRs (DYS19, DYS389I, DYS389II, DYS390, DYS391, DYS392, DYS393, DYS437, DYS438, DYS439, DYS448, DYS456, DYS458, DYS635, and Y_GATA_H4) using Arlequin Software v3.5[38]. Reduced dimensionality spatial representation of the populations was performed based on R_ST_ values using multi-dimensional scaling (MDS) with IBM SPSS Statistics for Windows, Version 23.0 (IBM Corp., Armonk, NY, USA). Heatmaps were generated using R_ST_ and F_ST_ values using the R program V3.4.1 platform [39] with the help of a ggplot2 module.

#### 2.4.2 Phylogenetic analysis

A neighbour-joining phylogenetic tree was constructed for the Qiang population and the reference populations based on a distance matrix of F_ST_ using the Mega7 software [40]. Y-DNA Haplogroup Predictor NEVGEN (http://www.nevgen.org) was used to predict Y-SNP haplogroups from Y-STR haplotypes. Any haplotypes which had null alleles or duplicated variants in the Qiang population were excluded from the analysis.

#### 2.4.3 The median-joining network

To define the genetic relationships among Qiang population individuals, we used the stepwise mutation model and Median Joining-Maximum Parsimony algorithm by using the program Network 5 as described at the Fluxus Engineering website (http://www.fluxus-engineering.com), and the weighting criteria for Y-STRs. Any haplotypes which had null alleles or duplicated variants in the Qiang population were excluded from the analysis.

### 2.5 Population admixture and evolutionary history reconstruction

#### 2.5.1 Principal component analysis and model-based ADMIXTURE

We conducted PCA analysis for 1158 individuals from 105 modern and ancient populations using smartpca of EIGENSOFT v.6.1.4 [41]. Ancient reference East Asians were projected onto the basic background of modern East Asians with the additional parameters (numoutlieriter: 0 and lsqproject: YES). We merged our genome-wide SNPs of 20 Qiang individuals with 1986 reference individuals to conduct a model-based ADMIXTURE analysis. We used PLINK v.1.9 [42] to prune the raw dataset using the following parameters (--indep-pairwise 200 25 0.4) and ran ADMIXTURE (v.1.3.0) [43] with 10-fold cross-validation and predefined ancestral sources. In-house script and PLINK v.1.9 [42] were used to calculate the pairwise F_ST_ genetic distances between Qiang and other reference populations.

#### 2.5.2 Allele sharing and admixture signatures in f-statistics

We estimated the shared genetic drift between Qiang people and other modern and ancient populations using the *qp3Pop* in ADMIXTOOLS in the form of *f_3_*(Qiang, reference; Mbuti) and estimated the admixture evidence using *qp3Pop* in ADMIXTOOLS [44,45] in the form of *f_3_*(source1, source2; Qiang). Additionally, we calculated *f_4_*-values in the form of *f_4_*(reference1. reference2; Qiang, Mbuti) and *f_4_*(reference1, Qiang; reference2, Mbuti) to test the asymmetrical/symmetrical relationship between Qiang people and others. We used qpAdm to estimate admixture proportion and ancestral sources of Qiang people and other neighbours. Both three-way and two-way admixture models were used here. And we used nine populations (Mbuti, Ust_Ishim, Kostenki14, Papuan, Australian, Mixe, MA1, Jehai and Tianyuan) as the outlier groups. To further explore the phylogenetic framework of the Qiang people, we used the *qpGraph* program implemented in the ADIXTOOLS 2 package [45,46] to test all possible evolutionary models and choose the best models based on the Z-scores and Minimum likelihood.

#### 2.5.3 Ancestral source composition and admixture dates inferred from chromosome painting

We phased on modern genomes in the merged dataset using SHAPEIT v2 [47] and used the major parameters as the following settings (--burn 10 --prune 10 --main 30). The phased haplotypes and refined-ibd.17Jan20.102 [48] were used to calculate the pairwise IBD between Qiang and modern reference populations. FineStructure v4.1.1 [49], ChromoPainter v2 and ChromoCombine v2 [49] were used to explore the fine-scale population structure and relationship of redefined genetic homogeneous populations. We finally used GLOBETROTTER to identify, date and describe admixture events.

## 3 RESULTS

### 3.1 Validation of the developmental 30-marker Y-STR amplification system

For sensitivity, we separately amplified control DNA 9948 and 007 with different concentrations (1.0 ng/μL, 0.5 ng/μL, 0.25 ng/μL, 0.125 ng/μL, 0.1 ng/μL, 0.05 ng/μL). Results showed that even as low as 0.125 ng of DNA template yielded complete amplification of AGCU Database Y30 kit loci (**Supplementary Figure 2**). When we reduced the DNA template concentration to 0.05ng, 80% of loci were not detected. So, this kit can effectively detect samples with a sensitivity of 0.125ng and hence AGCU Database Y30 kit demonstrated a high level of sensitivity.

For accuracy and consistency, we used different thermocyclers (GeneAmp^®^ PCR system 9700, Veriti^™^ Thermal Cycler, 2720 Thermal Cycler, and ProFlex^™^ PCR System (Applied Biosystems, CA, USA)) for the amplification of DNA templates from blood, saliva, filter paper, and FTA card. After amplification using the AGCU Database Y30 kit, the amplicons were detected through ABI3130XL, ABI3500, and ABI3730XL (Applied Biosystems, Waltham, MA, USA). There is no significant difference in amplification efficiency, the genotyping results are complete, clear and stable. There are no obvious non-specific peaks were seen with the AGCU Database Y30 kit. The genotyping results of the AGCU Database Y30 kit are almost the same with AmpFISTR^®^Yfiler plus^®^ on overlapping STRs (**Supplementary Figure 3**), which indicates that the kit has good accuracy and consistency.

For identity, we extracted DNA from blood, saliva and hair samples from the same source and amplified using the AGCU Database Y30 kit. The results showed that the DNA typing results of blood, saliva, and hair samples from the same source are all identical (**Supplementary Figure 4**), which means that the kit can precisely determine the genotype.

To check the stability of the AGCU Database Y30 kit, the components of the kit were repeatedly frozen and thawed 10 times and PCR amplification was performed on the final status (10^th^). Capillary electrophoresis results did not show any significant difference in amplification efficiency between different carrier samples (FTA card and filter paper) and the control which shows us the stability of this kit.

Generally, biological samples before recovering from crime scenes have been encountered in the natural environment which facilitates such agents which can cause inhibition in the process of polymerase chain reaction. The intermingling of such agents with biological samples can lead us to the failure of the amplification process and we cannot get a full profile. The most common inhibitors which frequently cause problems are from biological materials carrying themselves like hematin from blood and humic acid from the soil. Humic acid will bind to template DNA and hematin will alter Taq polymerase. To check the tolerance level of the AGCU Database Y30 kit, we have added six different inhibitors as mentioned in the M&M section. Typing results showed that a complete profile was generated when we used 40 and 50μg/μL of humic acid, 20 and 30μM of haemoglobin, 500, 750μM of EDTA, 0.5 and 0.75μM of Ca^2+^, 25 and 50μM of hematin and 4, 8μM of indigo. Hence this kit has shown a high tolerance level.

For specificity, we amplified 1-2.5 ng of genomic DNA from the human male, cat, sheep, chicken, duck, rabbit, mice, pig, E. coli and human female using the AGCU Database Y30 kit. We did not observe any non-specific DNA amplification which showed highly species-specificity (**Supplementary Figure 5**). We also amplified 20 unrelated female individuals and did not observe any peak after capillary electrophoresis, which means that the multiplex assay was gender specific.

Mixtures of DNA samples are frequent in forensic casework, mainly in sexual assault cases. We mixed control 9948 (male DNA) and control 9947A (female DNA) at various ratios of 1:1, 1:4, 1:9, 1:19, 1:29; which were amplified in triplicate. All mixtures accurately detected the male fraction (**Supplementary Figure 6A**). For the male-to-male mixture samples, control DNA 9948 and control DNA 007 were used to prepare mixtures with the ratio of 1:1, 1:4, 1:9, 1:19 and 1:29. These were amplified in triplicate. At the mixture ratio of DNA 007 is 1:19, the genotyping of each STR present in the AGCU Database Y30 kit was detected correctly (**Supplementary Figure 6B**).

### 3.2 PCR environment test results

#### 3.2.1 Reaction volume and primer concentration

We have used two different reaction volumes of 25 μL and 10 μL for amplification. Genotyping results showed us that there were no allele dropout or non-specific peaks and there was not any significant difference in peak heights for both the reaction volumes. When we used different primer concentrations (1X and 2X) for the amplification process, the genotyping results were consistent and there was no allele loss and non-specific peaks (**Supplementary Figure 7**).

#### 3.2.2 Annealing temperature and cycle numbers

Denaturation, annealing, and extension are the three vital steps for a successful PCR reaction. Annealing is the key step in PCR reaction which can determine the fate of its success or failure. Every amplification kit has a specific PCR annealing temperature which can fluctuate from one thermo-cycler to another. Therefore, it is important to check the tolerance level of the amplification kit to annealing temperature variations. We have used different annealing temperatures with variable cycle numbers and found that at 63 °C annealing temperature with 29 amplification cycles, the genotyping results were unstable and few allele peaks were lower than 100 (**Supplementary Figure 8**). At 59°C annealing temperature, with 29 amplification cycles, we got a perfect electropherogram with good peak heights (**Supplementary Figure 9**). So, 59°C annealing temperature, with 29 amplification cycles is the ideal annealing temperature and cycle number for the AGCU Y30 database kit.

### 3.3 Population studies and forensic parameters

#### 3.3.1 Allelic frequencies and forensic parameters

We successfully genotyped 30 Y-STRs in 514 males in the Qiang population residing in the Sichuan Province of China. 474 haplotypes were detected in the Qiang population. Haplotype data has already been made accessible via the Y-chromosome Haplotype Reference Database (YHRD) under accession number YA004683 (**Supplementary Table 1**). The gene or locus diversity (GD), allele frequencies and the number of observed alleles for 26 single-copy STR loci and 2 multi-copy STRs are summarized in **Supplementary Table 2**. Allele frequencies ranged from 0.0019 to 0.8326. Among single-copy STRs, the DYS481 showed the highest diversity of 0.8416 with thirteen different alleles, and the DYS391 was the least diverse locus 0.2856 with five different alleles. The multi-copy Y-STRs DYS385a/b and DYS527 showed more combinations of alleles when compared with single-copy Y-STRs. Gene diversities were 0.9324 and 0.9284 with 57 and 42 different allele combinations for DYS385a/b and DYS527, respectively. To assess the slow to medium mutating nature of the AGCU Y30 database kit, we evaluated the haplotype resolution with seventeen different combinations (Table 1) including the “minimal haplotype 9 Y-STRs”, “the extended 11 Y-STRs loci”, “PowerPlex Y12 STRs loci”, “Y-filer Plus 17 STRs” and so on till 30 Y-STRs. A total of 474 haplotypes were observed in 30 Y-STRs with haplotype diversity (HD) of 0.9996 and discriminatory capacity (DC) of 0.9222. Among 474 haplotypes, 85.79% haplotypes were unique. When we reduced the number of STRs to Yfiler 17 STRs, a total of 438 haplotypes were observed with HD value 0.9991 and DC value 0.8521. Among these 438 haplotypes, 76.26% haplotypes were unique. Multi-copy Y-STRs (DYS385a/b, DYS527) displayed 211 haplotypes with a HD value of 0.9729 and DC value 0.4105. Among these 211 haplotypes, 26.85% haplotypes were unique. This kit showed stronger discrimination power which means it can serve both purposes i) individual search and ii) familial search.

#### 3.3.2 Phylogenetic analyses and population comparisons

To reinforce the knowledge of the ethnohistorical records of Qiang and other Chinese populations, we used two different methods (pairwise R_ST_ and F_ST_ genetic distances) based on their similarity with *a priori* expectations. *F_ST_* is a standardized variance of haplotype frequency and assumes genetic drift as being the agent that differentiates populations. *R_ST_* is a standardized variance of haplotype size and takes into account both drift and mutation as causes of population differentiation, assuming a stepwise mutation model [50]. The genetic distances (*R_ST_* and *F_ST_*) between Qiang and other Chinese integrated populations are listed in **Supplementary Table 3~4**. Tibetan population from Amdo, China showed the closest genetic distance (0.108) with the Qiang population followed by the Manchu population (0.1205) from Jilin, China. On another hand, Yao population from Liannan, China showed maximum genetic distance (0.4629) followed by the Korean population (0.3851) from Yanbian, China among reference populations. According to F_ST_ genetic distance, Tujia population (0.0005) from Youyang, Chongqing, China showed the closest genetic distance which was followed by Han population (0.0006) from Anshan, China while Tuva population (0.0257) from Kanas, China showed the greatest genetic distance which was followed by Manchu population (0.0207) from Jilin, China. Phylogenetic relationships among the Qiang population and the reference populations were assessed using MDS analysis based on R_ST_ distances derived from the Y-STR data (**Supplementary Figure 10**). We have observed loosely three bounded clusters and the Qiang population was placed with the Tibetan population on the right side of the plot. Uighur, Mosuo, Salar, Yi and Kazakh populations clustered in the middle, while Han, Tujia, Maio, Gin and other populations clustered to the left side of the plot. In NJ tree, Qiang population was also placed along with Tibetan, Tuva, Manchu and Daur populations (**Supplementary Figure 11**). Genetic distances, MDS plot, and the phylogenetic tree showed that genetic affinity among studied populations was consistent with linguistic, ethnic, and geographical classifications.

#### 3.3.3 Ancestry information of Qiang ethnic groups using Y-STRs

Ancestry informative markers (AIMs) play an important role in genealogy. So, we used NEVGEN software to predict haplogroups from STRs. There were four haplogroups (D, J, O and R) that accounted for 87% while just two haplogroups (D and O) accounted for 75% of the Qiang population (**Figure 2**). Haplogroup J accounts for 4% of the Qiang population from China. This haplogroup J is predominately found in the Arabian Peninsula. The origin of this haplogroup is from the Middle Eastern area known as the Fertile Crescent, comprising Palestine, Jordon, Syria, Lebanon, and Iraq around 42.9KYA [51]. Merchants from Arabian Peninsula brought this genetic marker along with Silk Road to East Asia [52]. Haplogroup J was only observed in several groups from the extreme northwest of China which support the Silk Road spread phenomena. Haplogroup R originated from the north of Asia about 27 KYA (ISOGG, 2017). It is the most frequent haplogroup in Europe and Russia and, in some parts, it is at a frequency of 80%. Some researchers believe that one of its branches originated in the Kurgan culture [53]. In the Qiang population, the frequency of Haplogroup R was 8%. Previous studies [54–57] indicate that paternal haplogroup D originated in Central Asia. According to Hammer et al [54], haplogroup D originated between Tibet and the Altai mountains. He also speculated that multiple waves of human migrations swept into Eastern Eurasia. Haplogroup D, which is widely distributed across East Asia including most of the Tibeto-Burman, Tai-Kadai and Hmong-Mien-speaking populations. This haplogroup was predicted to be 36% in the Qiang population. According to Zhong et al [58], haplogroup D is present 2.49% in Han Chinese males. Frequencies of this haplogroup seem to be higher than average in the northern and western regions of China (8.9% in Shaanxi Han, 5.9% in Gansu Han, 4.4% in Yunnan Han, 3.7% in Guangxi Han, 3.3% in Hunan Han, and 3.2% in Sichuan Han). According to Yan et al [59], frequency of haplogroup D in the Han population from Eastern China (Jiangsu, Zhejiang, Shanghai, and Anhui) is 1.94%. The Tibetan population is thought to contain an admixture of two ancient populations represented by two major East Asian Y chromosome lineages, the O and D haplogroups [60]. Haplogroup O is the most common haplogroup in China and is prevalent throughout East and Southeast Asia. The frequency of haplogroup O in the Chinese Han populations ranges from 29.7% (Han from Guangxi) [61] to 74.3% (Han from Changting, Fujian) [62]. Haplogroup O is the most commonly found haplogroup in most East Asian populations such as approximately 40% in Manchu, Koreans and Vietnamese [54]; 34% in Filipino males [63]; 55% in Malaysian males [64]; 44% in Tibetan males [65]; 25% in Indonesian males [66] and 20% in Japanese male [54]. In Qiang’s population, its frequency was predicted to be approximately 39%. Haplogroup C has a very wide distribution and might represent one of the earliest settlements in East Asia. Haplogroup C was at low frequency in the Qiang population.

**Figure 2.**
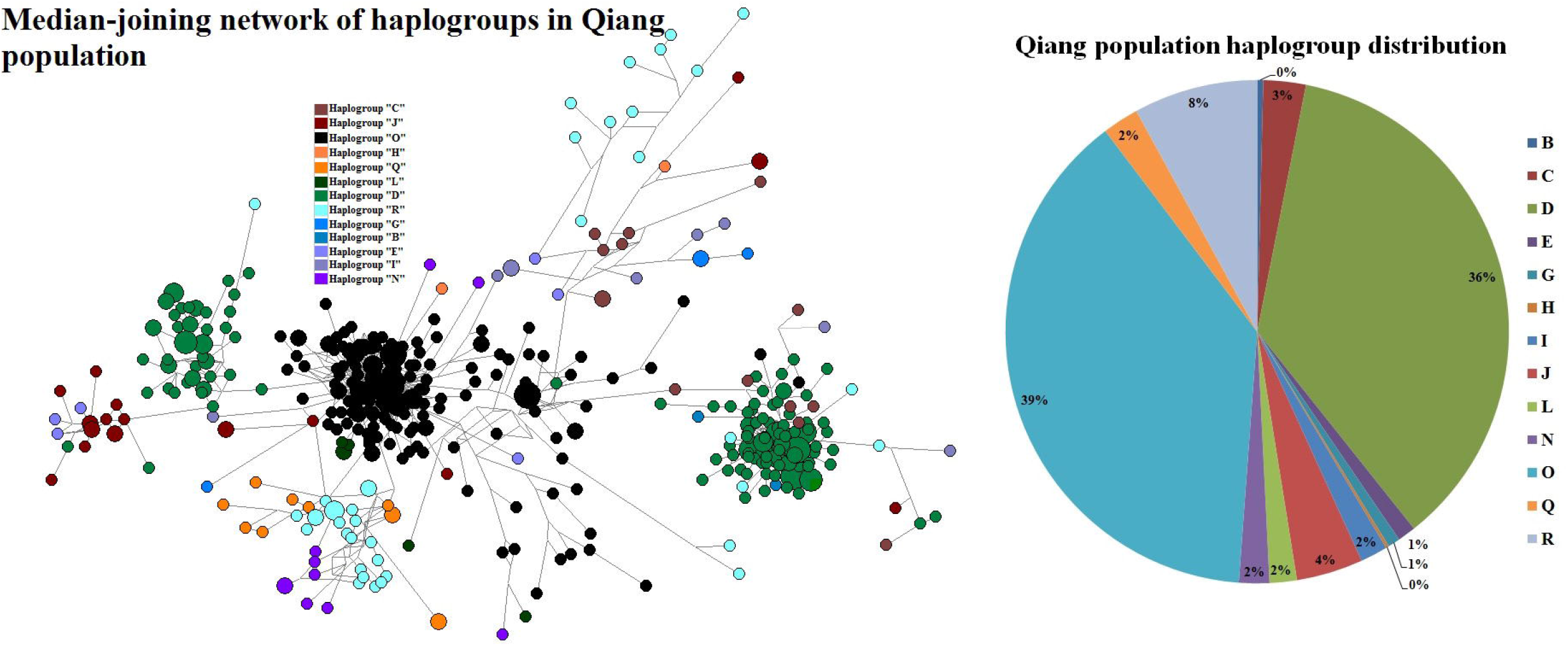
The Median-joining network of the predicted haplogroups in Qiang people and haplogroup frequency distribution.

### 3.4 Fine-scale admixture history and evolutionary frameworks of Qiang people

#### 3.4.1 Overview of the genetic structure of Qiang people inferred from genome-wide SNP data

To further formally identify, describe and date admixture events and characterize the detailed genetic landscape of Qiang people, we collected population data of 600K SNPs in 20 Qiang individuals from the Human Origins array and merged it with 1609 modern genomes and 377 ancient genomes from 59 eastern Eurasian populations. We firstly conducted the PCA analysis focused on the Chinese modern and ancient populations, in which ancient populations were projected onto the modern genetic landscape. We replicated five genetic lines respectively with geographically restricted language families (**Figure 3**). Ancient southern East Asians from Guangxi and Fujian clustered closely with Austronesian, Tai-Kadai and Hmong-Mien people related to modern northern East Asians. Ancient northern East Asians from the Yellow River Basin grouped closely to northern Hans and their neighbours, and ancient northeastern Asians showed a close relationship with modern Tungusic people. Qiang samples were localized closely with modern Tibetans but some of them with Han Chinese, which showed the existence of admixture events supporting the theory that Tibetans and Hans were the ancestral populations of Qiang people. These Qiang people with close genetic affinity with Hans were regarded as the outliers in the following formal tests to provide a better landscape of population admixture and evolutionary history of the indigenous Qiang people.

**Figure 3.**
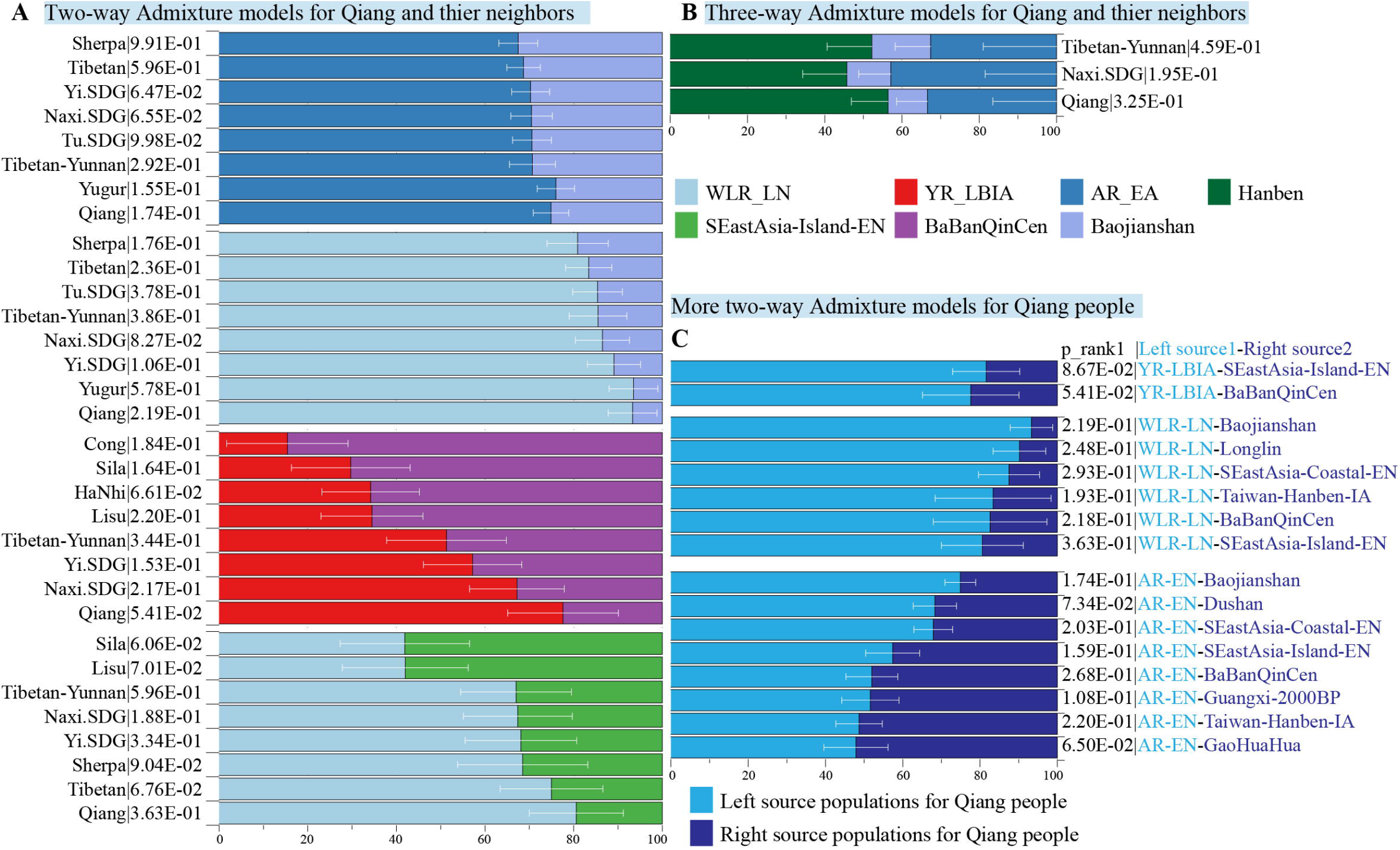
The principal component analysis results among 802 East Asian individuals from 76 modern and ancient populations. Each modern population was colored-coded by language family and ancient populations were color-coded by shape.

The ancestry composition and their corresponding admixture proportion of Qiang people were analysed using model-based ADMIXTURE software (**Figure 4**). We found 7.5% ancestries were related to Hmong-Mien-speaking Hmong, 8.3% were related to Austroasiatic-speaking Htin, 69.6% were related to middle Neolithic Miaozigou people and 0.146% ancestries maximized in Austronesian-speaking Atayal. Tibetans and ancient Nepalese people shared most ancestry composition of Qiang’s gene pool, suggesting their close genetic relationship with Yellow River Basin farmers. Additional genetic input from southern China related to ancient Guangxi and Fujian people also played an important role in the formation of the Sichuan Qiang people. To further dissect the genetic difference between geographically diverse Qiang people, we merged our data with 1875 individuals from 231 Eurasian populations and conducted a model-based ADMIXTURE analysis based on the bested fitted model with seven ancestral sources (**Figure 5**). We found Danba Qiang and Daofu Qiang people shared a similar pattern of admixture history and derived most of their ancestry from the ancient Nepalese population (83.4~83%). Similar to the first fitted model, Atayal-related southern East Asians also contributed to the Qiang people’s gene pool (15.8~16.2%). Pairwise F_ST_ genetic distances can provide direct evidence for genetic differentiation through inter-populations and intra-populations heterozygosities. We first identified the closest genetic relationship between Danba Qiang and Daofu Qiang (F_ST_: 0.0043, **Supplementary Table 5**). We also identified a close genetic relationship between Chuanxi Qiang and ancient Mongolian and Xiongnu. The estimated F_ST_ values between Qiang people and modern East Asian reference populations showed that Qiang had the least genetic distances with Naxi and Yi, followed by Hans. Compared to ancient populations, we found Qiang people had a closer genetic relationship with middle and upper Yellow River Basin people. We also used ‘outgroup-*f_3_*-statistics’ to measure the genetic affinity of geographically different populations and we found two geographically different populations (Daofu and Danba) shared a close genetic relationship, followed by Tibetans, Nepal ancients and Neolithic Lajia people (**Figure 6 and Supplementary Table 6**). Generally, the patterns between Danba Qiang and their reference populations are consistent with the pattern observed focused on Daofu Qiang. Finally, we calculated the ‘pairwise outgroup-*f_3’_* value among the meta-Qiang and their reference populations (**Figure 1**). We identified consistent patterns of genetic affinity between Qiang and reference modern and ancient East Asians in these two indexes. Modern Qiang people showed a close genetic relationship with Tibetans and ancient Yellow River Basin farmers who are geographically close to each other.

**Figure 4.**
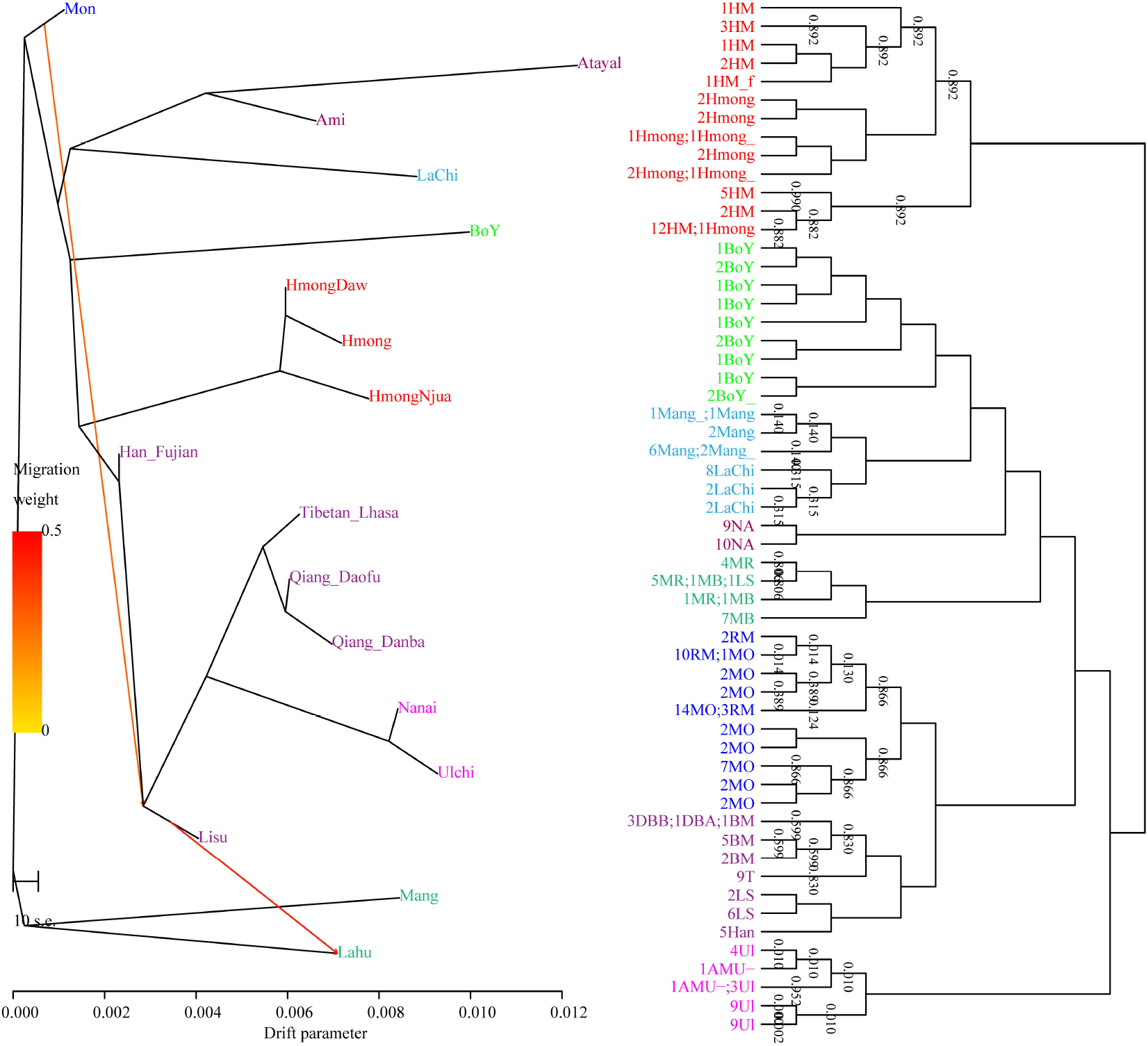
Model-based ADMIXTURE results among 1788 East Asian individuals from 133 modern and ancient populations. Each population was visualized with equal width for better presentation for the population label, which is not associated with population size. K equal to six has the least cross-validation error.

**Figure 5.**
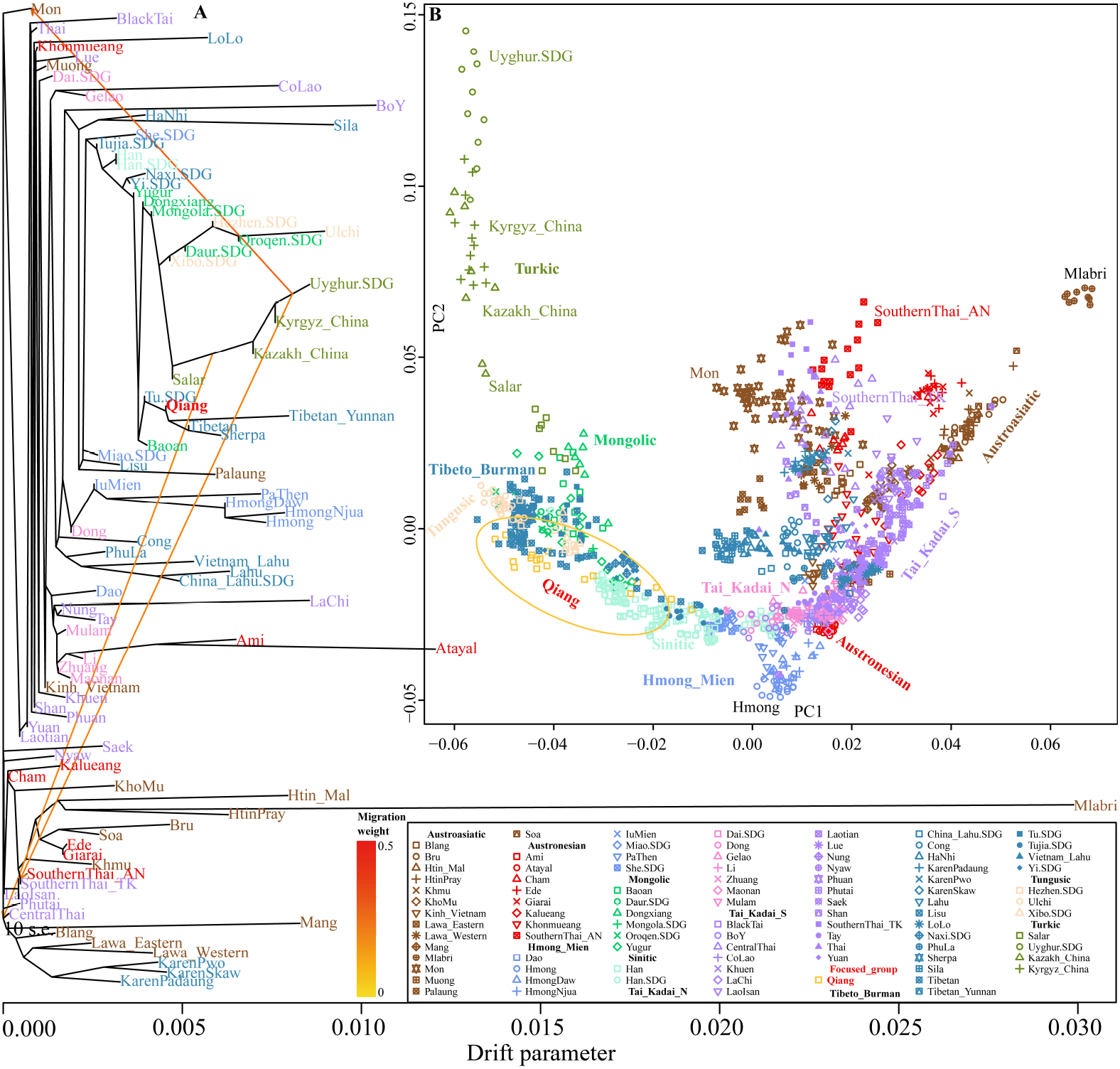
Ancestry composition of 233 modern and ancient Eurasian populations based on the model-based ADMIXTURE model. We used the merged HO and Illumina datasets here. K equal to six has the least cross-validation error.

**Figure 6.**
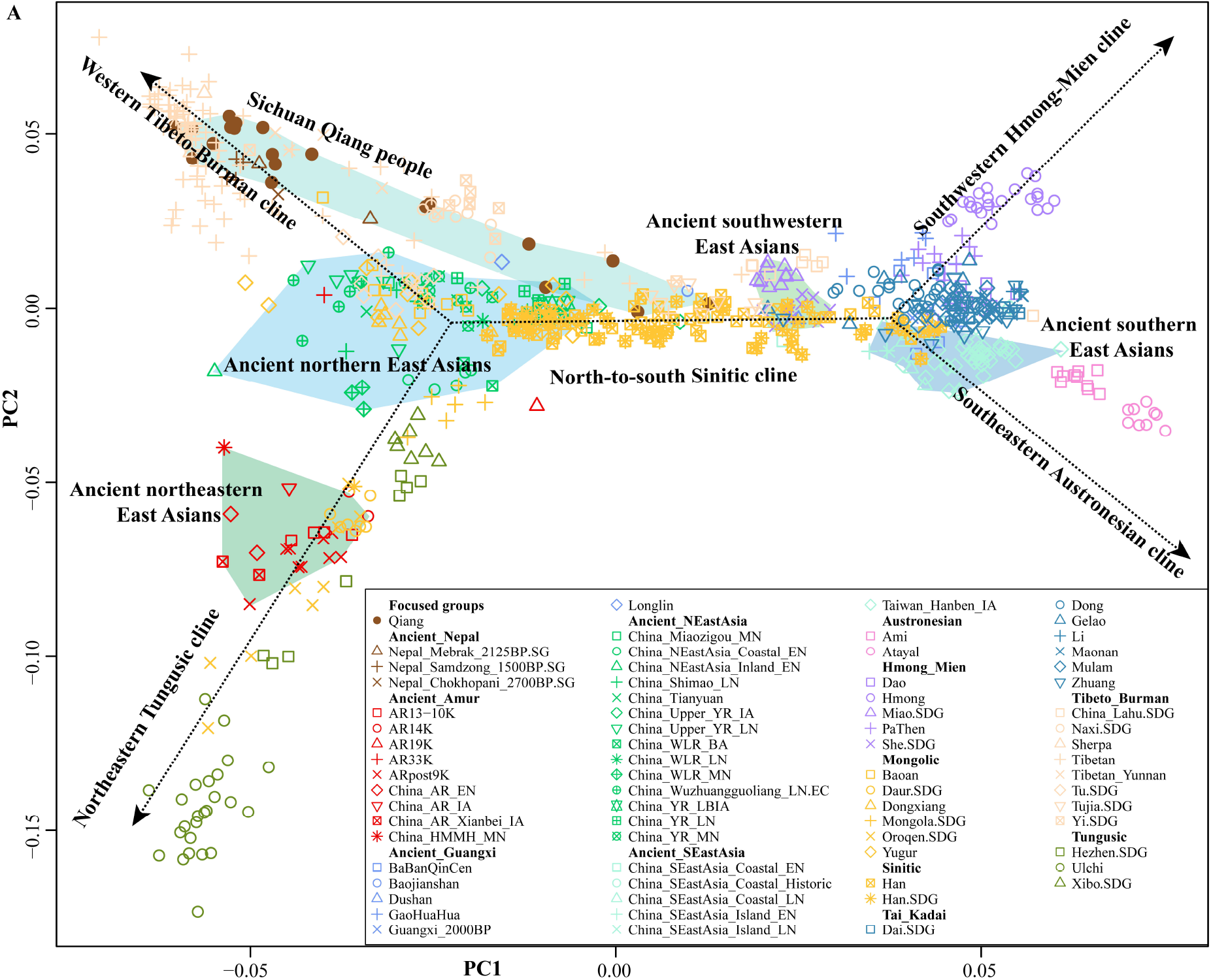
Shared genetic drift between Danba and Daofu Qiang people and Eurasian reference populations. Bar showed the genetic affinity between Qiang people and ancient Eurasian populations. Circle size and color showed the genetic affinity between Daofu Qiang and modern reference populations.

#### 3.4.2 Formal testing for the admixture process of Qiang people

We calculated ‘admixture-*f_3_*-statistics’ in the form of *f_3_*(source1, source2; Qiang) to formally explore the formation process of Qiang populations, in which the statistically negative values (Z-scores less than −3) indicated that the targeted populations were a mixed product with the ancestral surrogate related to source1 and source2. Most Tibeto-Burman-speaking populations, such as highland Tibetan and Yunnan Lahu, showed no statistically significant signals in the admixture *f_3_*-statistics. Here, we found many source pairs with admixed evidence for Qiang people with one source from northern East Asians (Tibetan and Yellow River Basin farmers) and the other one related to the southern East Asians (Tai-Kadai and Austronesian speakers, **Supplementary Table 7**). Additionally, we conducted the *f_4_*-statistics in the form of *f_4_*(Reference1, reference2; Qiang, Mbuti) to test the asymmetrical or symmetrical sharing ancestry signals between Qiang people and modern and ancient East Asians. Among modern reference populations, we found that Qiang people share most alleles with Yi, Naxi, Tibetans and Hans compared with other reference populations. When compared with ancient East Asians from southern East Asia, or Siberia, Qiang shares more alleles with middle and late Neolithic people from Yellow River Basin (**Figure 7**). Finally, we performed *f_4_*-statistics in the form of *f_4_*(reference1, Qiang; reference2, Mbuti) and we found many signals generating negative values with reference2 for modern Sino-Tibetan people or ancient Yellow River farmers, suggesting additional gene flow from these populations contributed to modern Qiang people. Generally, our *f_3_/f_4_* analyses showed Qiang people were a mixed population with ancestral sources derived from their northern and southern neighbours.

**Figure 7.**
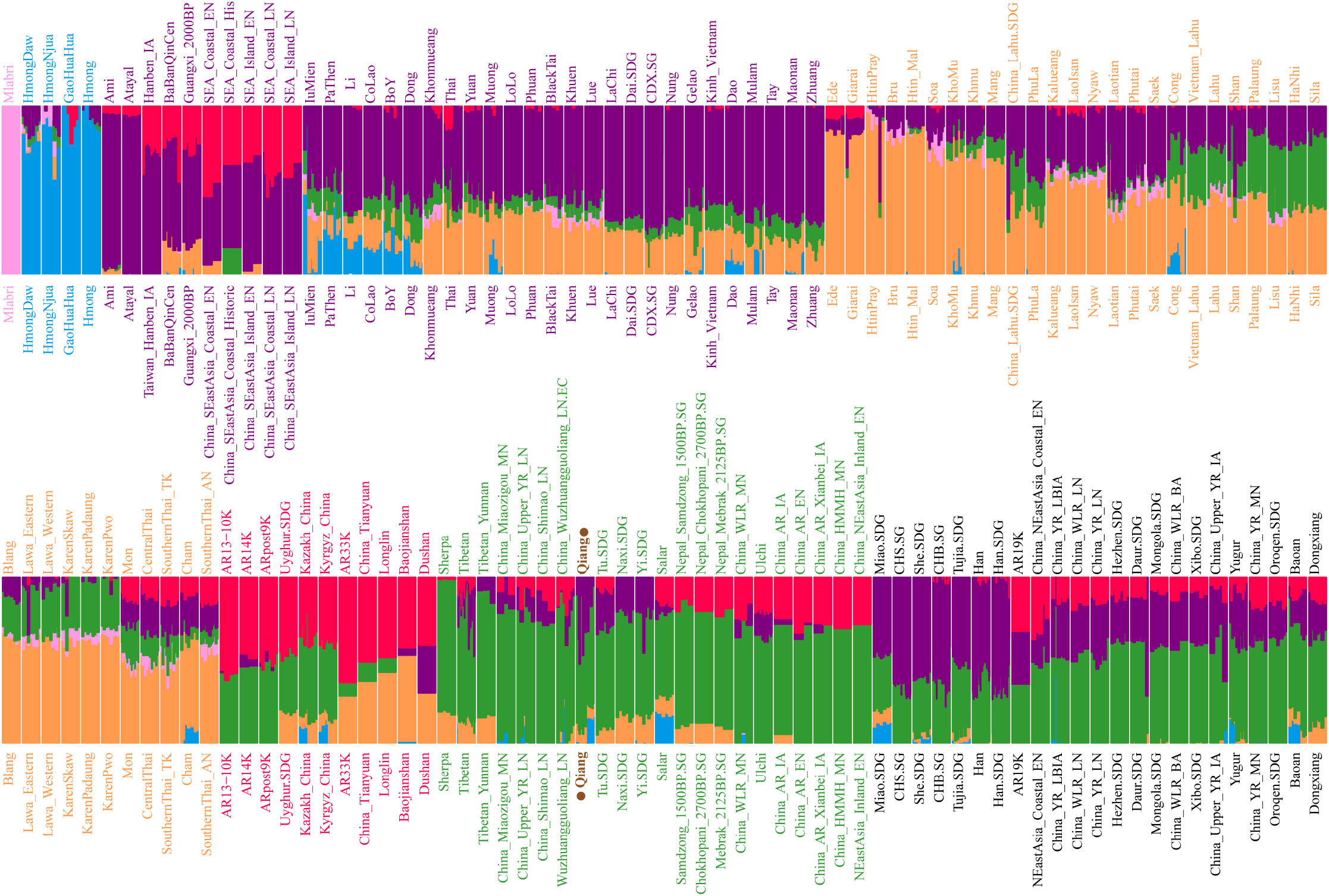
Results from *f_4_*-statistics showed the genetic affinity between Qiang and northern East Asians. The statistically significant population pairs were marked with stars and the red color showed the Qiang showed a close relationship with the right populations compared to the bottom populations. Blue color with negative values showed a close genetic relationship with bottom populations compared with the right populations.

We formally estimated the ancestry source composition and admixture coefficient of Sichuan Qiang people and their geographically close populations. Considering the identified two-way admixture signatures from northern and southern China, we first conducted the two-way admixture qpAdm models to portray their ancestry admixture processes using Early Neolithic people from Amur River Basin (China_AR_EN), Neolithic people from the West Liao River Basina (China_WLR_LN) and Bronze/Iron Age populations from the Yellow River Basin (China_YR_LBIA) as the northern sources and using Neolithic people from Fujian and Guangxi and historic people from Guangxi as the potential southern sources. Our spatiotemporal analysis focused on the admixture process. We found that Qiang and other northern Tibeto-Burman-speaking populations (Tibetan and Sherpa) derived their primary ancestry from the ancient northern East Asians in the predefined two-way admixture models. Southern Tibeto-Burman people harboured more ancestry related to the southern East Asians, such as Sila. It was modelled as an admixture result of 0.297 ancestries related to Yellow River farmer ancestry and the remaining ancestry appeared to be the BaBanQinCen, which is consistent with the admixture patterns observed in Cong people (**Figure 8A**). To further validate whether both inland and coastal southern East Asians contributed to the formation of the Sichuan Qiang people, we conducted a three-way admixture model using China_AR_EN as the northern source, Baojianshan as the inland southern source and Hanben_IA as the coastal source. We found Qiang people could be modelled as an admixture of the 0.0564±0.095 ancestry from China_AR_EN, 0.102±0.080 ancestry from Baojianshan and 0.334±0.165 ancestry from Hanben (**Figure 8B**). The admixture models could well fit the formation of the Naxi people and Yunnan Tibetans. Finally, we also tested other models with different north-south source pairs for Qiang people, and we confirmed the major ancestry of Qiang people descended from ancient northern East Asians (**Figure 8C**).

**Figure 8.**
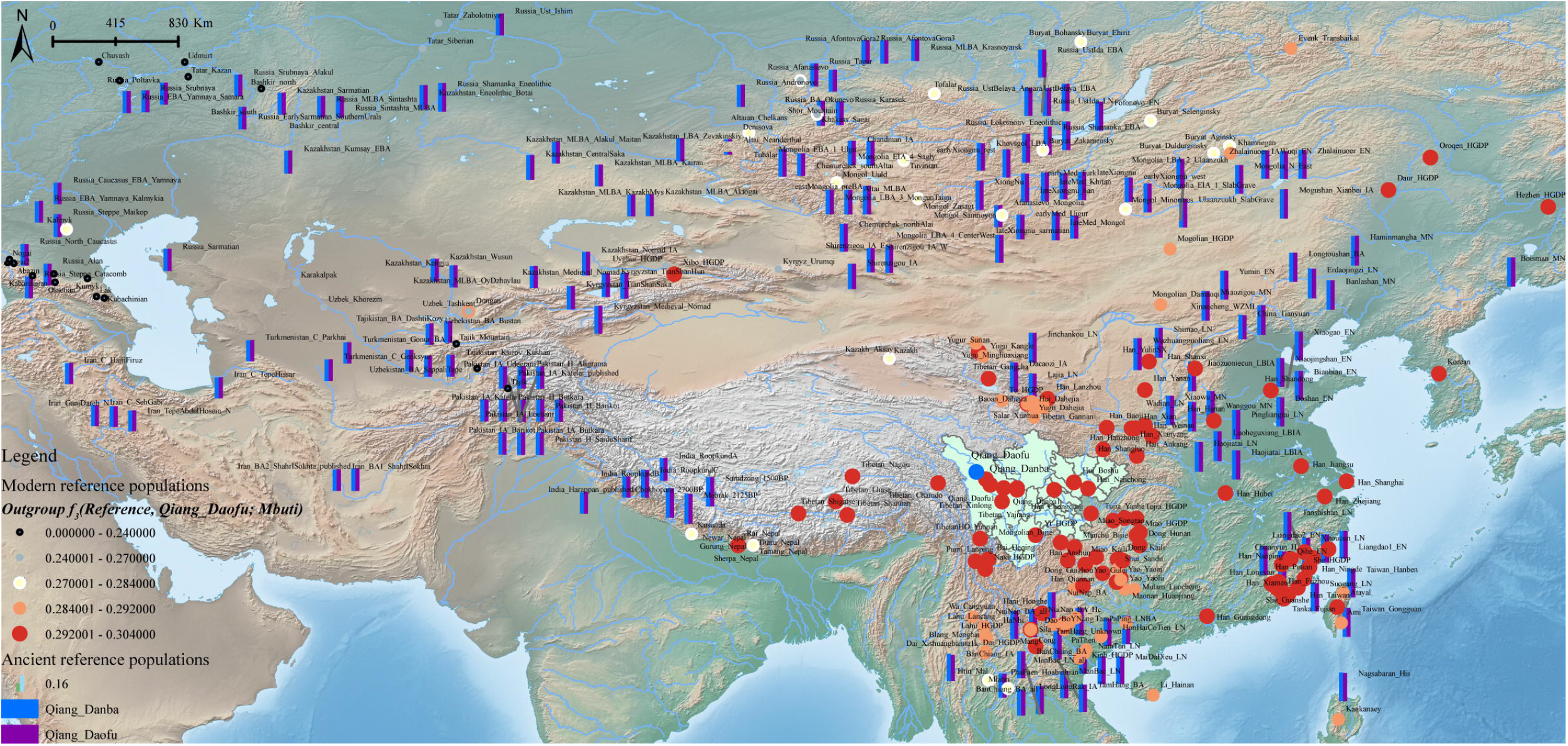
Results of the qpAdm-based models show the formation of modern Tibeto-Burman-speaking populations. Error bar denoted the standard error and different colors showed different ancestral sources.

The length and number of ancestral chromosome fragments document more traces of the human population in the process of evolutionary history under different driving forces, including natural selection, mutation, drift and recombination. We followingly conducted a population genetic analysis based on phased haplotypes (**Figure 9**). Shared IBD showed that Qiang people had the closest genetic relationship with modern Naxi, Yi and Tibetans. PCA based on the co-ancestry matrix showed a similar cluster pattern, which is also consistent with the identified genetic affinity via allele sharing in the *f*-statistics. Fine-scale population substructure based on the genetic similarity of re-classified homogeneous populations also confirmed the genetic affinity between Qiang and other lowland Tibeto-Burman speakers. Finally, we painted the targeted chromosomes of Qiang people using both northern and southern East Asians as the potential ancestral sources (surrogate populations) using the Chromo-Painterbased copy vector. We identified the one-date-multiway admixture model as the best model guesses for the Sichuan Qiang people. GLOBETROTTER-based admixture dates showed that Qiang people underwent multiple admixture events in the historic times with the surrounding populations, a northern source related to the Tibetans and southern sources related to the southwestern indigenes. We finally reconstructed the phylogenetic relationship among Chinese populations (**Figure 10**). Consistent with the population clustering patterns among the PCA results, the reconstructed phylogenetic relationship showed that Qiang people clustered closely with northern East Asians related to Tibetan, Sherpa and Tu people, suggesting their common population history.

## Discussion

The origin and admixture history of the Qiang people has been a mystery in China. Culturally-documented evidence supported the common origins of Tibetan and Qiang people, some also suggested a close relationship between Qiang and Han people. However, no population genetic study focused on the fine-scale population history of Qiang populations has been reported. In this study, firstly, we developed and comprehensively validated a 30-plex Y-STR multiplex assay and found that it is robust enough for forensic genetics applications. Secondly, we genotyped the Qiang population for use as forensic genetic reference data. We identified high diversity and high efficacy for forensic application. Population genetic analysis based on the Y-STR data showed a close relationship between Qiang and Tibetan people. Thirdly, we made a comprehensive population genomic analysis focused on the genetic structure and admixture history of modern Qiang people.

Findings from PCA and ADMIXTURE analyses showed Qiang people clustered closely to the geographically close Yi, Tu and Tibetans. Previous genetic analysis showed Tibetan populations harboured the pre-Neolithic ancestry related to the early Asian and Neolithic ancestry from northern Chinese millet farmers [67]. Geographically diverse Tibetans also showed differentiated population structure, such as Tibetan in the Tibetan-Yi corridor showed more ancestry related to southern East Asians [8,11]. We also identified different genetic structures between Qiang and highland Tibetans. Qiang people in Sichuan Province showed a mixed landscape with major ancestry from Yellow River farmers and minor ancestry related to southern East Asians (Dai and Atayal), suggesting that Qiang possessed more genetic influence from southern Chinese populations. Our ADMIXTURE and qpAdm results consistently supported the most ancestry of Qiang people derived from northern East Asians, which was consistent with the common origin of Qiang, Tibetan and Han Chinese populations from the Yellow River Basin in North China. The direct evidence for the close relationship between Qiang and Neolithic millet farmers was obtained from the identified closest relationship with Qiang and Late Neolithic Lajia and Jinchankou people from the Ganqing region via shared genetic drift testing. We also identified the close genetic relationship between Qiang and historic Xiongnu and Mongolian, which was consistent with these ancient populations resided in the wide regions of northwest China contributed to the genetic materials into modern Qiang people. We also noted that the genetic history of only two geographically different Qiang populations was explored, we believed that more genome-wide SNP data from geographically different regions can provide deeper and more comprehensive insights into the complex landscape of the Qiang people.

## Conclusion

This study first performed a developmental validation work focused on the AGCU Database Y30 kit and population genetic analyses based on the Y-STR diversity and genome-wide SNP variation. The study showed that the Y30 kit was a robust assay for forensic casework and can be used as a powerful tool for forensic identification and population genetics research. Population genetic analysis based on the Y-STR and genome-wide SNP data showed that Qiang people had the closest genetic affinity with geographically close Tibeto-Burman-speaking Naxi, Yi and Tibetan populations. Further evidence from shared ancestry in *f*-statistics and qpAdm-based admixture models showed Qiang people derived most of their ancestry from Neolithic Yellow River millet farmers and the remaining ancestry was derived from the Neolithic-to-historic Guangxi people. This is consistent with the reconstructed population evolutionary of modern Tibetan people based on the modern and ancient people. Furthermore, we also identified obvious recent admixture models in the one-date-multiway model based on the phasing modern chromosome painting. This study supported the view that Qiang people primarily originated from the Yellow River Basin related to millet farmers and underwent continuous gene flow events from southern East Asians from ancient to the historic time period.

## Acknowledgments

This study was supported by Princess Nourah bint Abdulrahman University Researchers supporting project number (PNURSP2022R318) Princess Nourah bint Abdulrahman University Riyadh Saudi Arabia. GLH was supported by Project funded by China Postdoctoral Science Foundation (2021M691879). We thank Prof. Wibhu Kutanan in Khon Kaen University, Prof. Mark Stoneking and Dr. Dang Liu in Max Planck Institute for Evolutionary Anthropology for sharing genome-wide SNP data from Vietnam, Thailand and Laos.

## Data Availability

The Genome-wide variation data were collected from the public dataset. The new-generated Y-STR data has been submitted to the YHRD database with the access number YA004683.

## Figure Legends

**Figure 9. Phylogenetic clustering patterns based on the sharing alleles and the reconstructed haplotypes**. (A) TreeMix-based clustering patterns with two migration events. (B) Fine-structure results showed the close relationship between Qiang and Tibetan people. The same color denotes the same populations.

Figure 10. TreeMix-based phylogenetic relationship between Qiang and other East Asians and PCA results of these batch of populations.

